## Supplementary figures and images for "Human Papillomavirus Genomes Associate with Active Host Chromatin during Persistent Viral Infection"

### Supplementary Figures

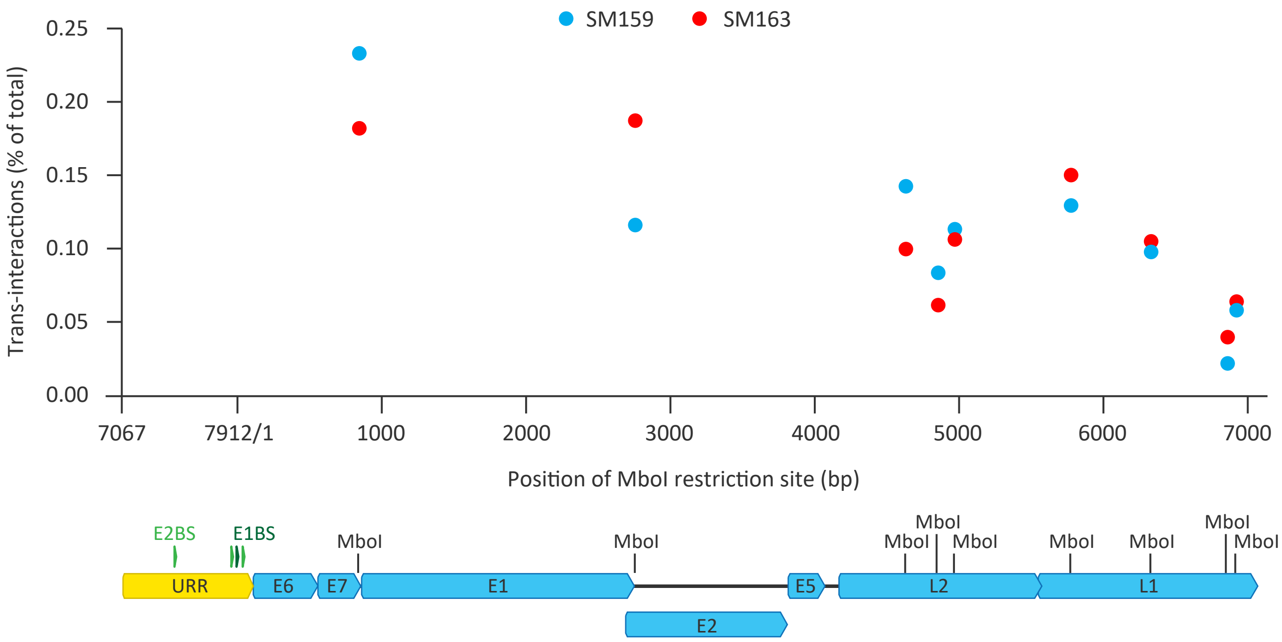

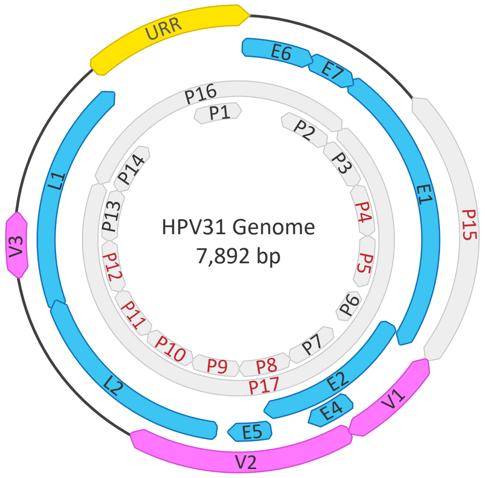

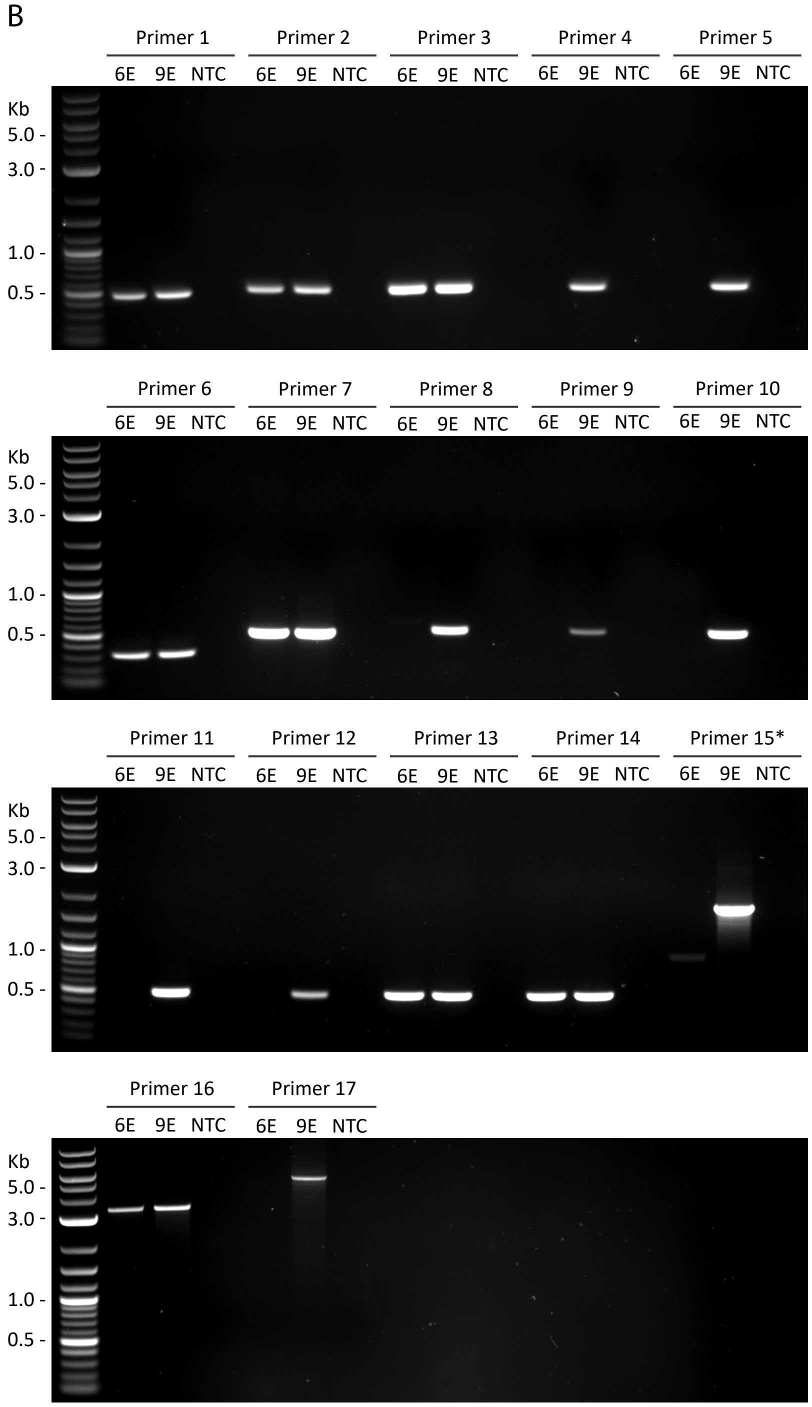

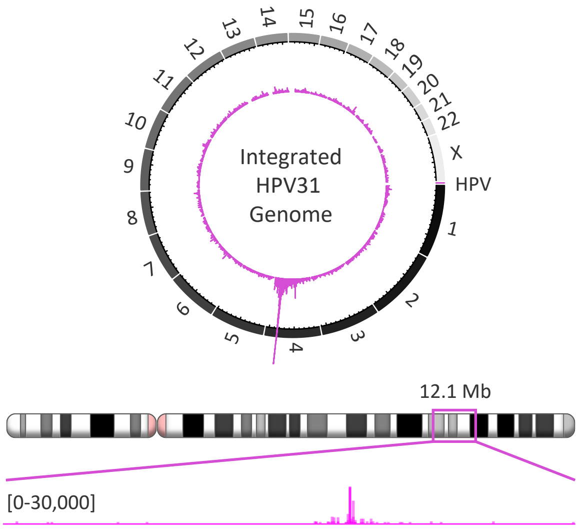
